# Long non-coding RNAs in wild wheat progenitors

**DOI:** 10.1101/301804

**Authors:** Alice Pieri, Mario Enrico Pè, Edoardo Bertolini

## Abstract

*Triticum urartu* and *Aegilops tauschii* are the diploid progenitors of the hexaploid *Triticum aestivum* (A^u^A^u^BBDD), donors of the A^u^ and D genome respectively. In this work we investigate the long noncoding RNAs (lncRNAs) component of the genomes of these two wild wheat relatives. Sixty-eight RNA-seq libraries generated from several organs and conditions were retrieved from public databases. We annotated and characterized 14,515 *T. urartu* and 20,908 *Ae. tauschii bona*-*fide* lncRNA transcripts that show features similar to those of other plant and animal counterparts. Thousands of lncRNAs were found significantly modulated in different organs and exhibited organ specific expression, with a predominant accumulation in the spike, fostering the hypothesis of their crucial role in reproductive organs. Most of the organ-specific lncRNAs were found associated with transposable elements (TEs), indicating the possible role of TEs in lncRNA origin, differentiation and function. The majority of *T. urartu* and *Ae. tauschii* lncRNAs appear to be species-specific; nevertheless, we found some lncRNAs conserved between the two wheat progenitors, highlighting the presence and conservation of exonic splicing enhancers sites in multi-exon conserved lncRNAs. In addition, we found cases of lncRNA conservation and their *cis* regulatory regions spanning the wheat pre-domestication and post-domestication period. Altogether, these results represent the first comprehensive genome-wide encyclopedia of lncRNAs in wild wheat relatives, and they provide clues as to the hidden regulatory pathway mediated by long noncoding RNAs in these largely unexplored wheat progenitors.

## 1. Introduction

Bread wheat (*Triticum aestivum*, 2n=6×=42, A^u^A^u^BBDD) is the result of the fusion of three diploid sub-genomes that occurred over a temporal scale of 10 MYA (Marcussen et al. 2014). *Triticum urartu* and *Aegilops tauschii* were proposed as the donors of the A^u^ and D genome respectively, whereas the source of the B genome is still debated (Kilian et al. 2007). In 2013 the draft genomes of both *T. urartu* and *Ae. tauschii* were published (Jia et al. 2013; Ling et al. 2013), and recently the improved *Ae. tauschii* reference genome (Aet v4.0) was released (Luo et al. 2017). Overall, diploid wheat together with the whole-genome sequences of the tetraploid progenitor wild emmer (*T. turgidum ssp. dicoccoides*, 2n = 4x = 28, A^u^A^u^BB) (Avni et al. 2017) and of bread wheat (Marcussen et al. 2014; Clavijo et al. 2017) represent a precious resource to decipher the complexity of wheat biology, its genome structure and its evolution. Moreover, wild wheats serve as an extraordinary source of exotic genes and alleles, in particular for abiotic and biotic stress tolerance, which are useful for future genomics-assisted crop breeding programs (Feuillet et al. 2008).

However, in order to fully exploit the wheat wild relative gene pool in future plant breeding schemes, it is crucial to deepen our knowledge regarding the molecular mechanisms controlling their gene action, especially in response to stresses. In recent years, several studies have reported the intricate regulatory network activated in response to abiotic stimuli (Pearce et al. 2015). However, the relevant fraction of the genome encoding for noncoding RNAs that emerged as a major component of the eukaryotic transcriptome (Ben Amor et al. 2008) has been mostly neglected and remains largely unexplored. In particular, long noncoding RNAs (lncRNAs) are mostly transcribed from the intergenic space and characterized by transcript units longer than 200 nt with biochemical features that resemble those of messenger RNAs. LncRNAs contribute to modulating gene expression in a broad range of mechanisms and in a multitude of pathways (Ulitsky 2016), including plant developmental programs and abiotic/biotic stress tolerance (Chekanova 2015).

Few functional studies have clarified the epigenetic involvement of lncRNAs in controlling gene expression through chromatin modification mechanisms in plants. In *Arabidopsis*, the polycomb-dependent repression of the *FLOWERING LOCUS C*, during the vernalization process, is mediated by *COLD ASSISTED INTRONIC NONCODING RNA* and *COLD INDUCED ANTISENSE INTERGENICRNA* (Ietswaart et al. 2012); the auxin polar transporter *PINOID* expression in root is controlled by the *AUXIN REGULATED PROMOTER LOOP RNA* (Ariel et al. 2014). In rice, the lncRNA *LONG DAY SPECIFIC MALE FERTILITY ASSOCIATED* RNA (*LDMAR*) is required for normal pollen development and its transcription is epigenetically regulated through the photoperiod (Ding et al. 2012).

The target mimicry mechanism is another well-known mechanism through which lncRNAs act. *INDUCED BY PHOSPHATE STARVATION*, involved in the phosphate uptake, interacts with miR-399 through a partial complementarity, thus preventing the cleavage of *PHOSPHATE 2*, the miR-399 target, which, in turn, negatively affects phosphate shoot content and its remobilization (Franco-Zorrilla et al. 2007). Despite these well-defined examples, the systematic characterization of lncRNA loci is still in its infancy, although in the past few years a number of rigorous genome-wide mapping lncRNA studies based on extensive RNA-sequencing have been published for various plant species, such as *Arabidopsis thaliana* (Liu et al. 2012; Wang et al. 2014), *Brachypodium distachyon* (De Quattro et al. 2017), rice (Zhang et al. 2014; Wang et al. 2016) (Shuai et al. 2014; Chen et al. 2015; Tian et al. 2016) and tomato (Zhu et al. 2015).

Here we present a comprehensive annotation of long noncoding RNAs in the wild wheat progenitors *T. urartu* and *Ae. tauschii* based on a large dataset comprising 68 public RNA-seq libraries generated from different plant organs. In total, 14,131 and 20,523 intergenic lncRNA transcripts were identified in the draft genomes of *T. urartu* and *Ae. tauschii*, respectively. Wild wheat lncRNAs showed shorter and a reduced number of exons compared to protein-coding genes. We observed that lncRNAs are characterized by a specific and dynamic expression pattern in most of the plant organs, and they predominantly accumulate in developing spikes, suggesting their potential functional role in reproductive organs. Our comparative analyses identified some lncRNA transcripts conserved between *T. urartu* and *Ae. tauschii* and, for those lncRNAs characterized by a multi-exon gene structure, we showed the conservation of exonic splice enhancers (ESEs) motifs. The interspecific conservation of lncRNA sequences and regulatory regions was maintained through the domestication process, from the wild diploid species to the wild tetraploid emmer, to cultivated polyploid wheat species. This finding is of particular interest because it highlights, for the first time, the evolutionary conservation in lncRNA sequence and structure between wild relatives and their polyploid product that merits further studies to assess whether lncRNAs might have played a role in wheat speciation and domestication.

## 2. Results

### A comprehensive catalogue of wild wheat relative long noncoding RNAs

To identify lncRNAs in the wild wheat progenitors, we retrieved data from sixty-eight public RNA-seq libraries from 7 organs (root, shoot, leaf, leaf in cold stress and control conditions, seedling and spike) in *T. urartu* and from 10 organs (root, shoot, leaf, seedling, seed, spike, pistil, sheath, stem, stamen) in *Ae. tauschii* (supplementary table S1).

All the computational analyses were conducted according to the protocol described in De Quattro et al. (De Quattro et al. 2018). Briefly, after the quality check of the raw reads and the removal of sequencing adapters, reads from each library were aligned against their own reference genome (see Material and Methods) through two mapping iteration steps using the aligner TopHat2 v2.0.9 (Trapnell et al. 2012). On average, 72% and 77% of the reads were mapped to the draft reference genome of *T. urartu* and *Ae. tauschii* respectively, giving proof of the quality of the draft assemblies. The transcriptome was assembled with Cufflinks v2.0.9 (Trapnell et al. 2010) and overall 176,863 and 186,966 transcripts were reconstructed in *T. urartu* transcripts and in *Ae. tauschii* respectively. These complete sets of transcripts were then used to identify high-confident lncRNAs on the basis of the current features of lncRNA biogenesis and functions (Quinn and Chang 2016; Ulitsky 2016). To distinguish lncRNA candidates from the coding transcripts, five sequential stringent filters were employed (supplementary fig. S1). More precisely: i) transcript sequences shorter than 200 nucleotides and with ORFs longer than 100 amino acids were discarded; ii) the remaining transcripts were aligned against the entire Pfam protein database to identify known protein domains; iii) quality, completeness and sequence similarity of potential ORFs were assessed using the program Coding Potential Calculator (CPC) (Kong et al. 2007); iv) housekeeping RNAs were then removed using a homology-based approach against a custom *Poaceae* housekeeping RNA database (see Materials and Methods); v) to reduce the complexity of the two datasets without losing low-abundance transcripts, we decided to retain only those lncRNAs having an overall expression of at least 10 read counts in total across the different organs, in accordance with the study from Gaiti *et al.* (Gaiti et al. 2015).

Overall, this computational approach led to a total of 14,515 *T. urartu* and 20,908 *Ae. tauschii bona*-*fide* lncRNAs. LncRNA isoforms within each species were resolved based on a sequencing identity of 95% using the program CD-HIT-EST (Li and Godzik 2006) resulting in a non-redundant set of 13,993 lncRNA loci in *T. urartu* (Tu-lncRNA) and 20,338 in *Ae. tauschii* (Aet-lncRNA) (supplementary file S1 and S2).

To further strengthen our lncRNA assembly, we compared the two lncRNA datasets to the *T. aestivum* EST databases NCBI (http://www.ncbi.nlm.nih.gov/). In addition, our Tu-lncRNA dataset was also compared with the full set of *T. urartu* noncoding transcripts *de novo* reconstructed by Krasileva et al. (Krasileva et al. 2013) (See Materials and Methods for details). Within the *T. aestivum* NCBI EST database 20,721 hits were found with *T. urartu* and 26,532 in *Ae. tauschii* (supplementary file S3 and S4). Moreover, the comparison between Tu-lncRNAs and *T. urartu* noncoding transcripts generated 1,574 hits (supplementary file S5). Altogether these results show that although the biological samples available in the ESTs databases do not perfectly match the organs present in our dataset, our *in silico* reconstruction of lncRNAs is generally confirmed by Sanger sequencing. It addition, it should be considered that the ESTs used for this analysis belong to *T. aestivum* and lncRNAs are transcripts characterized by low conservation among different species and by a temporal/tissue-specific transcription profile (Quinn and Chang 2016).

To further consolidate the Aet-lncRNA annotation we used the recently improved genome assembly (Aet v.4.0) (Luo et al. 2017) to map back the consensus transcripts derived from the draft genome version. Considering only perfect matches over the entire sequence, we retrieved 14,141 hits equally distributed on the seven chromosomes, of which 312 were duplicated (supplementary file S6 and supplementary fig. S2). The duplicated lncRNAs were found tandemly arrayed either on the same or different chromosomes, irregularly spaced and seldom inverted, in groups of 2, 3, 4, 5, 6, 7, 8, 9, 10, 11, 12, 13, 14, 15, 19. Notably, the lncRNA TCONS_00001910, duplicated six times, was found clustered in tandem in a 12 Kb region on chromosome 3 within the protein-coding gene *AET3Gv20726400* (a putative enhancer of mRNA-decapping protein) (supplementary fig. S2). We also found four lncRNAs duplicated more than 20 times across the entire genome (TCONS_00078769 [n=21]; TCONS_00031939 [n=22]; TCONS_00093600 [n=37]; TCONS_00143101 [n=47]). To address whether these duplication events could be due to a burst in TE activity, we masked the nucleotide sequences with a TE *Aegilops tauschii* custom library (see Materials and Methods). Out of 312 of the duplicated Aet-lncRNAs 147 (47%) resulted masked mainly with LTR and TIR elements. Specifically, the TCONS_00143101, duplicated 47 times, contained a TIR TE domain within its sequence.

The remaining lncRNAs, not perfectly mapped to the new reference genome, were located either on the 200 Mb of the super-scaffolds remained unassigned (Luo et al. 2017) or mapped onto the reference genome by allowing one mismatch.

### Genomic features of long noncoding RNAs

The main features of *bona fide* lncRNAs were compared with the predicted coding gene set of *T. urartu* and *Ae. tauschii*, considering length distribution, GC content and exon number (fig. 1). In agreement with the main characteristics described in the literature of plant lncRNAs (Liu et al. 2012; L. Li et al. 2014; Shuai et al. 2014; Wang et al. 2014; Zhang et al. 2014; Chen et al. 2015; Zhu et al. 2015; De Quattro et al. 2017), our data suggest that wild wheat lncRNAs are mostly mono-exonic (77% in *T. urartu* and 80 % in *Ae. tauschii)* or harbor fewer exons than protein coding genes. In fact, in both species, 94% of lncRNAs are characterized by 1 or 2 exons. We also found that lncRNAs are on average shorter than protein coding genes (465 nt vs. 861 nt in *T. urartu* and 464 nt vs. 987 nt in *Ae. tauschii*, p-value < 2.2e-16), but no significant differences in term of GC content was found between lncRNAs and coding genes.

**Figure 1.**
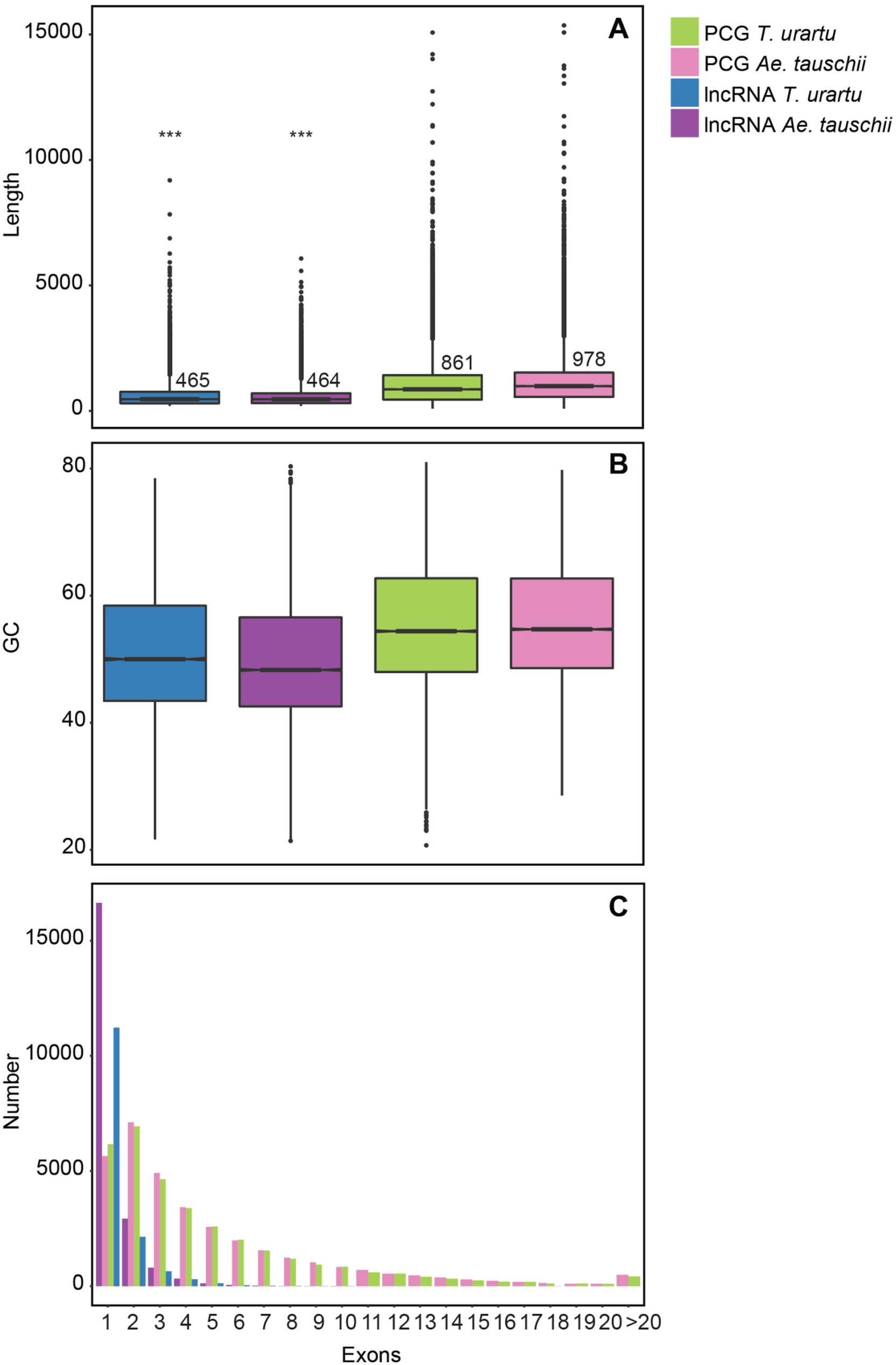
Structural characteristics of *T. urartu* and *Ae. tauschii* lncRNAs. Genomic features of the identified *T. urartu* and *Ae. tauschii* lncRNAs and reference protein coding genes. (A) length distribution (numbers indicate the median length), (B) GC content, (C) number of exons. *T. urartu* lncRNAs (blue), *Ae. tauschii* lncRNAs (violet), *T. urartu* protein coding genes (PCG) (green), *Ae. tauschii* protein coding genes (PCG) (pink).

To avoid bias arising from the complication of working with draft genome sequences, we focus our attention specifically on long intergenic noncoding RNAs (lincRNAs) which, according to their genomic location, do not overlap with exons or introns of protein-coding genes (Ulitsky and Bartel 2013). In addition, lincRNAs are the product of RNA polymerase II, holding the typical biochemical features of mRNAs. They are thus expected to be the most abundant class of lncRNAs in the poly(A)+ RNA-seq libraries (Ulitsky 2016). According to their genomic location, within the full set of lncRNAs herein identified, we classified 14,131 *T. urartu* and 20,523 *Ae. tauschii* lincRNAs (97% and 98% of the identified lncRNAs in the two species, respectively) that do not overlap with protein coding genes.

### Transposable element content of long noncoding RNAs

*T. urartu* and *Ae. tauschii* genomes have been shown to contain approximately 60% of transposable elements (TEs) sequences, with *Copia* and *Gypsy* long terminal repeat (LTR) retrotransposons as the most abundant families (Jia et al. 2013; Ling et al. 2013). In the new *Ae. tauschii* assembly (Zhao et al. 2017) TE accounts for 85.9% of the genome. In human and mouse, it has been shown that a large fraction of lincRNA exonic regions derive from transposable elements (Kelley and Rinn 2012; Kapusta et al. 2013), leading to the hypothesis that exonic TE could serve as functional RNA domains implicated in their function and evolution (Johnson and Guigó 2014). Despite these observations, there are few works that have studied the relationship between TEs and lncRNAs at the genome scale, especially in plants (De Quattro et al. 2017; Wang et al. 2017). We investigate the possibility that a lncRNA might be transcribed from an annotated TE locus and the presence of TE domains within the exonic sequences of a lncRNA (see Materials and Methods). We found that 1,376 Tu-lncRNAs and 4,156 Aet-lncRNAs overlap with the annotated TE coordinates, highlighting an overrepresentation of LTR *Gypsy* and *Copia*, followed by *CACTA* DNA transposons (fig. 2). Moreover, by masking the lncRNAs using a species-specific TE library (see materials and methods) we found that 27% of lncRNA exonic sequences were found to be associated with TEs domains in *T. urartu* and 19% in *Ae. tauschii*, and largely belonging to LTR retrotransposons (23% in *T. urartu* and 14% in *Ae. tauschii).* This result is in sharp contrast with vertebrates, where more than two thirds of lncRNAs contain an exon sequence of TE origin (Kapusta et al. 2013). This difference could be biased by the low coverage and incomplete annotations of the wild wheat genomes compared to the human, mouse and zebrafish genomes, the ones mostly characterized for lncRNAs.

**Figure 2.**
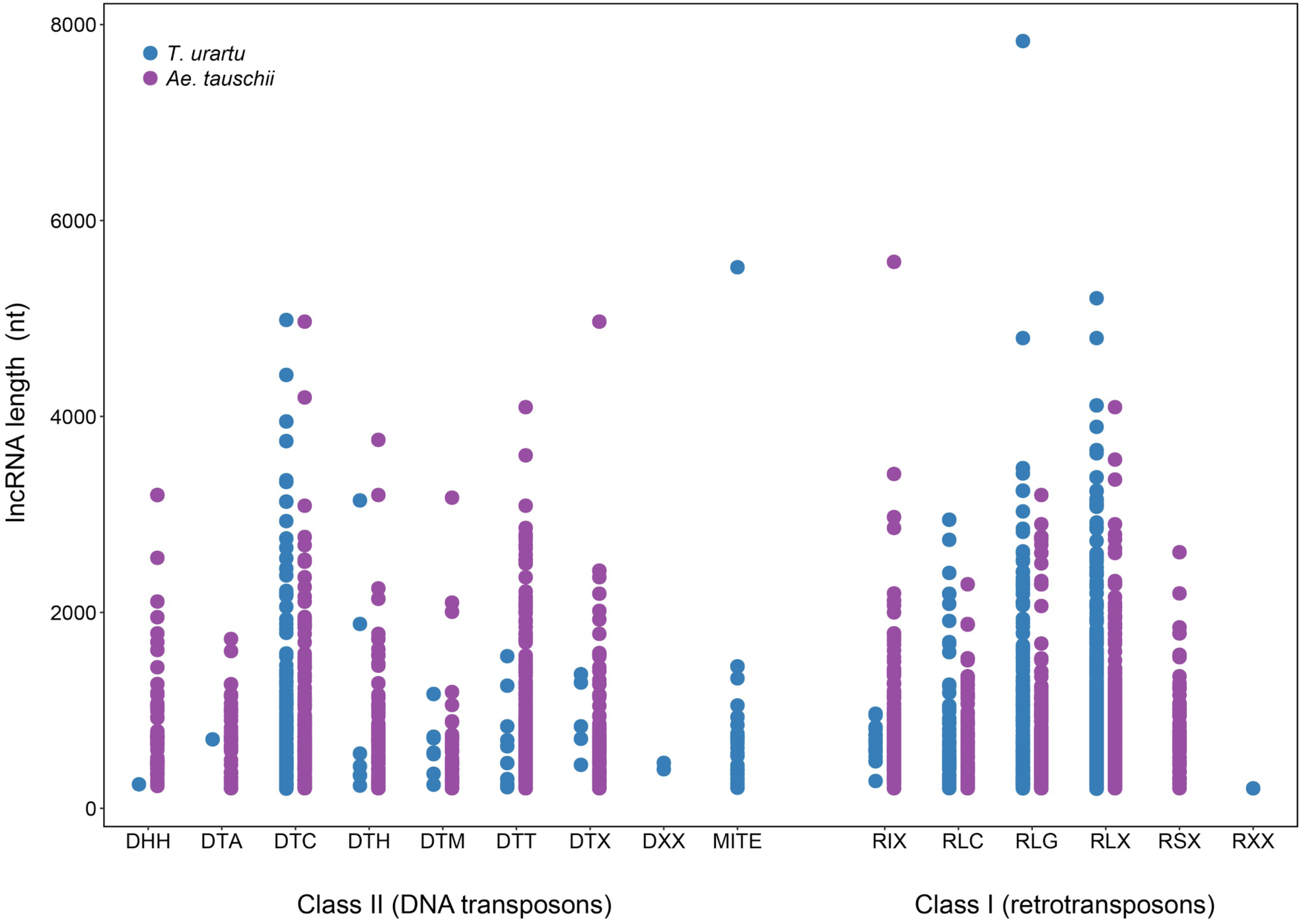
*T. urartu* and *Ae. tauschii* TEs associated with lncRNAs. Dot plot of the different TE classes enriched in lncRNAs. Dots represent lncRNAs transcribed form TE coordinates. TE code based on the classification system proposed by Wicker et al (Wicker et al. 2007).

### Conservation of lncRNAs between wild wheat progenitors

LncRNAs appear to be poorly conserved during evolution, implying a rapid evolution of the primary nucleotide sequence and structure, although a certain degree of synteny and conservation of the secondary RNA structure has been observed (Diederichs 2014). Using a homology-based approach, we first checked for the presence of sequence identity to assess the evolutionary conservation of lncRNAs between the wheat progenitors (*T. urartu* vs. *Ae. tauschii*) by aligning the query lncRNA with lncRNAs from other species (see materials and methods). Overall, 5,400 unique hits between *T. urartu* and *Ae. tauschii* were obtained. Most of these lncRNAs showed a conserved gene structure (65% mono-exonic and 7% with a multi-exon gene structure), while the remaining 28% were characterized by a non-conserved exon-intron structure, displaying sequence identity across the exons. Interestingly, within this last group, we found 200 mono-exonic lncRNAs where the sequence identity was spliced by an intronic region in one of the two wild wheat (fig. 3A). By imposing a sequence identity cutoff of at least 80%, we identified 164 lncRNA clusters of near perfect sequence identity spanning the entire length of the transcript (fig. 3 and supplementary file S7). Among the 356 lncRNAs characterized by a conserved multi-exon gene structure, we also looked for the presence of conserved exonic splicing enhancer (ESE) motifs, because these sequences appear to evolve at a lower rate than non-ESE sequences and are involved in splice-mediated lncRNA selection, at least in humans (Blencowe 2000; Schüler et al. 2014; Haerty and Ponting 2015). ESEs are known as purine rich hexamers that can be bound and activated by serine arginine-rich (SR) proteins, leading to the recognition of the splicing site (Schaal and Maniatis 1999; Blencowe 2000). Among the ESE candidates, we selected only the sequences proved to be very effective ESEs in plants (see materials and methods), obtaining 151 lncRNAs containing one or more ESEs. As reported in the supplementary Table S3, we found that most of the ESEs are in the same exon between *T. urartu* and *Ae. tauschii* and 109 out of the 151 ESE motifs have almost the same distance from the splicing site in the two wild species, confirming the trend seen in humans (Schüler et al. 2014; Haerty and Ponting 2015).

**Figure 3.**
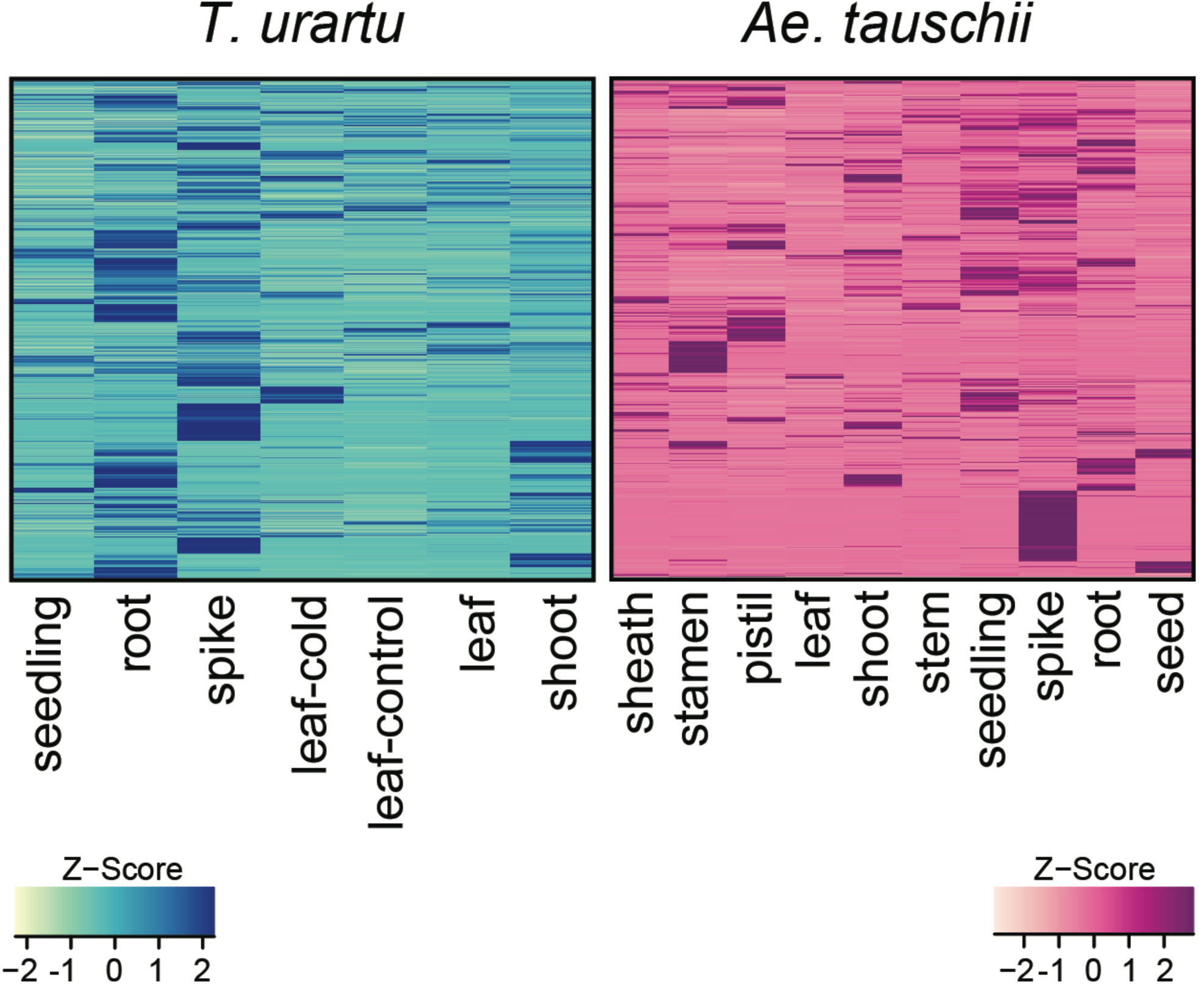
*T. urartu* and *Ae. tauschii* IncRNAs expression profile. Expression profiles of *T. urartu* and *Ae. tauschii* IncRNAs across different samples. Expression levels normalized in RPKM (Reads Per Kilobase Million) were rescaled in Z-Score to better compare trends across samples. Columns were hierarchically clustered (Ward method). Light colors indicate low expression levels, dark colors high expression. Each row represents data for one lncRNA.

### LncRNA conservation through the domestication of wheat

To provide insight into the conservation of lncRNAs through wheat evolution and domestication, we searched for lncRNA conservation patterns among the tetraploid emmer wheat and the hexaploid bread wheat, using a whole-genome alignmens approach (see Materials and Methods for details).

Tu-lncRNAs were compared with the genomic sequences of *T. aestivum* and *T. turgidum* ssp. *dicoccoides* genome A, whilst Aet-lncRNAs were compared against the genomic sequence of *T. aestivum* genome D. In accordance with wheat evolution, the degree of wild wheat lncRNA conservation drastically varied between the A and the D genomes, following the evolutionary distance. In particular, we found 2,074, 1,850 and 926 lncRNAs conserved between *Ae. tauschii* and *T. aestivum, T. urartu* and *T. turgidum* and *T. urartu* and *T. aestivum*, respectively (fig 3).

These data were further validated using all the wheat EST present in the NCBI EST database (see Materials and Methods) to verify whether or not this conservation reflects genomic conservation, but also if it concerns transcription activity. Notably, 823 and 536 lncRNAs, respectively in *T. urartu* and in *Ae. tauschii*, were supported by Sanger sequences in *T. aestivum* (supplementary file S8-S9).

In contrast to the lack of sequence conservation in the gene body, the *cis*-regulatory regions of lncRNAs seem to be more conserved (Amaral et al. 2016). To investigate this phenomenon we also achieved the conservation analysis of lncRNA promoters among the wheat species, bringing out the conservation of 2,022 and 3,581 regulatory regions in the A genomes respectively between *T. urartu vs.* bread wheat and *T. urartu vs.* emmer wheat. In addition 4,227 promoters were found conserved between the D genomes of *Ae. tauschii* and bread wheat.

It is of great interest that by comparing the A sub-genome of all three species, 557 lncRNA and 1,888 regulatory regions are highly conserved throughout the evolution and domestication process (supplementary file S10).

### LncRNA expression in different organs

The lncRNA expression level was examined to identify significant changes among the different organs taken into consideration in this work. *T. urartu* and *Ae. tauschii* lncRNAs were expressed at a low level across the samples with strong organ specificity (fig. 4). Based on the Shannon Entropy score (see materials and methods), we found that both wild wheats have a higher portion of organ-specific lncRNAs (supplementary fig. S3), in particular spike-specific lncRNAs (n=669 in *T. urartu* and n=2,273 in *Ae. tauschii*). This organ-specific expression was also found in root (n=221 in *T. urartu* and n=205 in *Ae. tauschii*), seed (n=355 in *Ae. tauschii)* and stamen (n=325 in *Ae. tauschii*). These organs are characterized by active growth and differentiation and often they are generally highly expressed as shown in the boxplots in fig. 5. In this work we also examined whether the expression of lncRNAs associated with TEs vary among the data set, detecting a dynamic expression profile of both class I and II TEs. Notably, both species showed a large number of TE-lncRNAs highly expressed in the spike and in particular in the floral-organs of *Ae. tauschii*, such as stamen and pistil (fig. 6).

**Figure 4.**
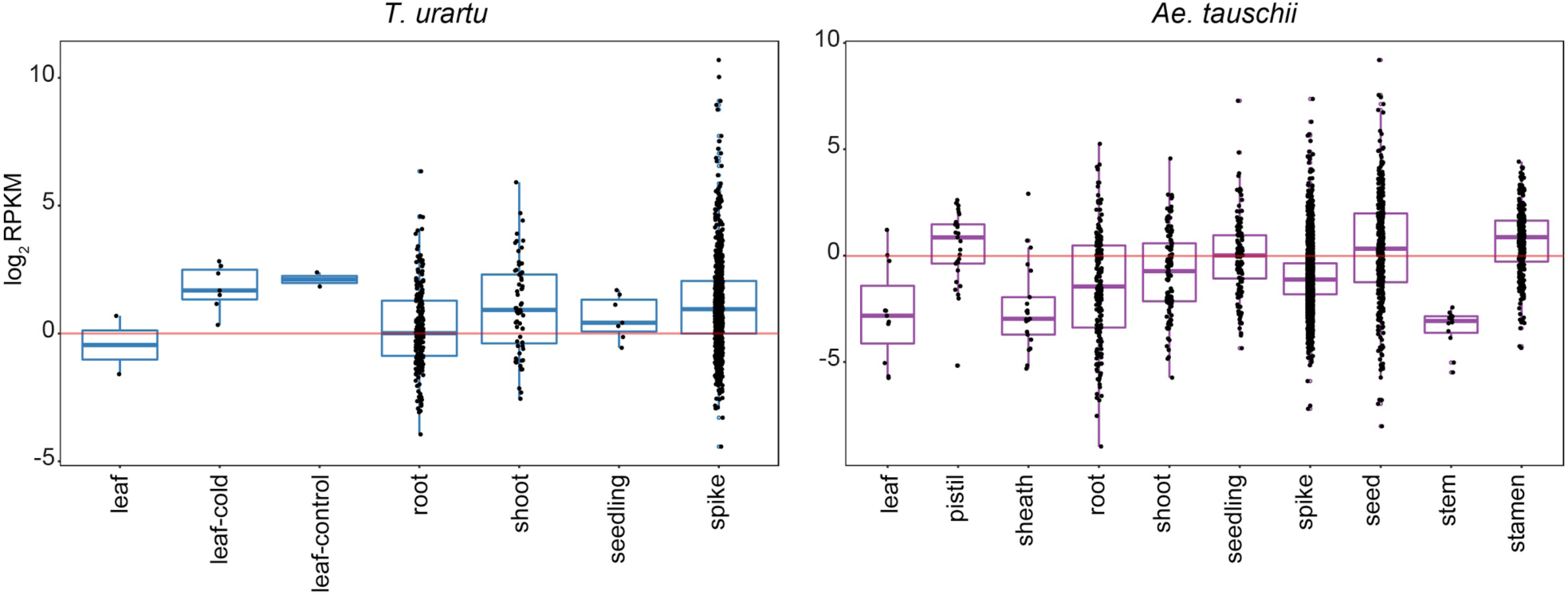
*T. urartu* and *Ae. tauschii* lncRNA organ-specific expression. LncRNAs characterized by organ-specific expression. The black dots represent the number of lncRNAs specific to each organ. The expression level is expressed in log2 RPKM (Reads Per Kilobase Million).

**Figure 5.**
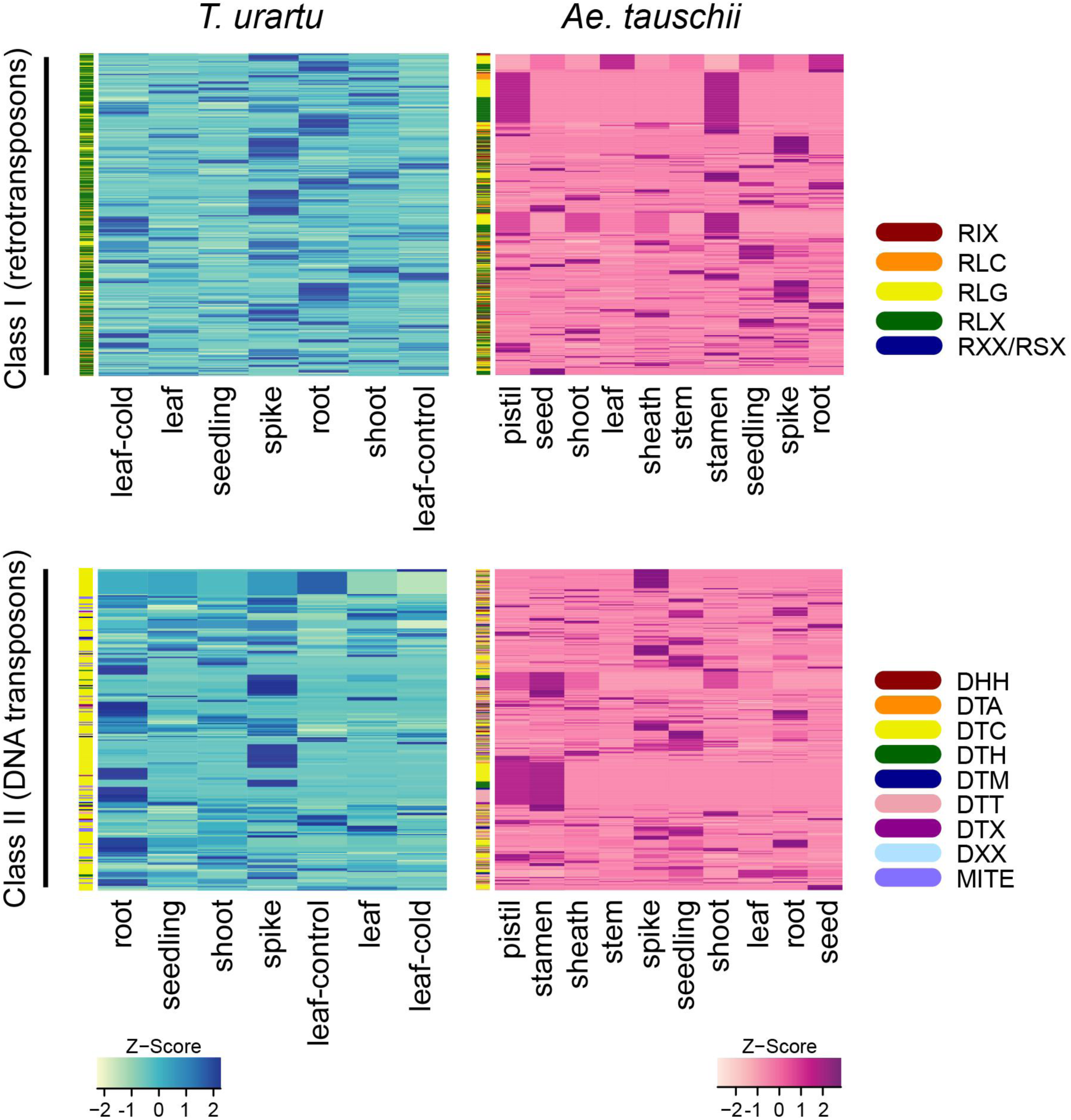
Expression profile of *T. urartu* and *Ae. tauschii* TEs associated with lncRNAs. *T. urartu* and *Ae. tauschii* expression profiles of lncRNAs associated with TEs across the different organs. Expression levels normalized in RPKM (Reads Per Kilobase Million) were rescaled in Z-Score to better compare trends across samples. Columns were hierarchically clustered (Ward method). Light colors indicate low expression levels, dark colors high expression. Each row represents data for one lncRNA.

**Figure 6.**
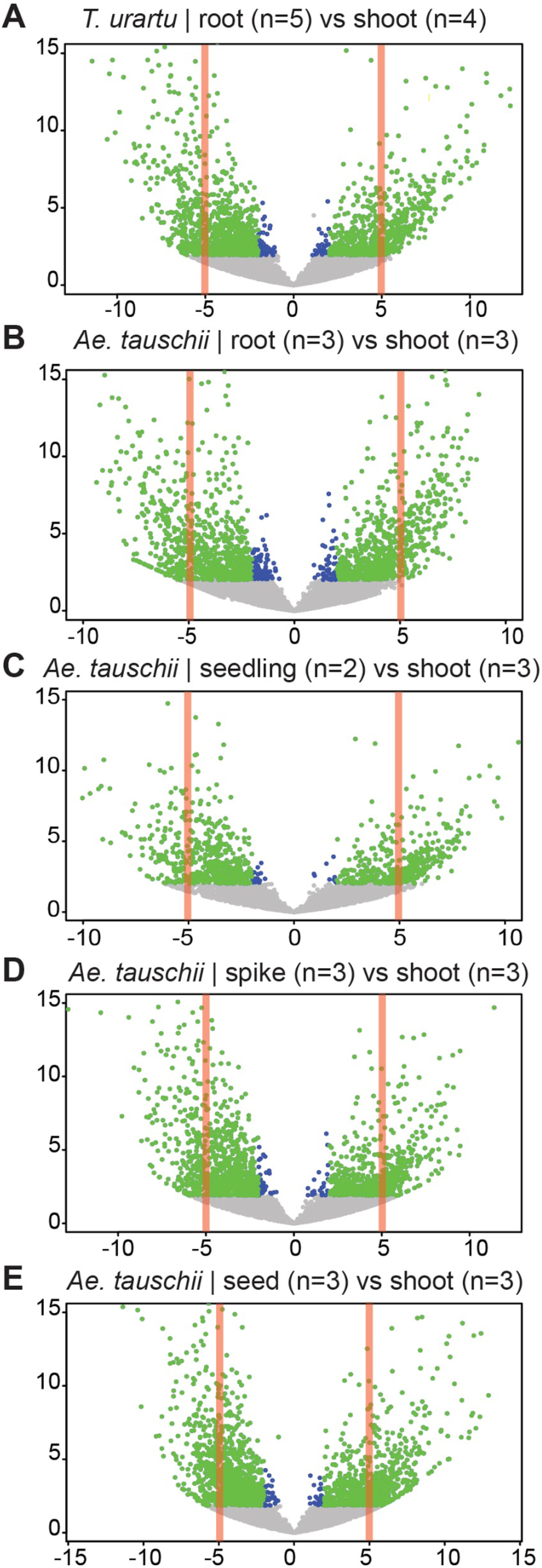
Differentially expressed IncRNAs. Volcano plots showing the differentially expressed (DE) IncRNAs in each pairwise comparison. (A) *T. urartu* root *vs.* shoot. (B) *Ae. tauschii* root *vs.* shoot. (C) *Ae. tauschii* seedling *vs.* shoot. (D) *Ae. tauschii* spike *vs.* shoot. (E) *Ae. tauschii* seed *vs.* shoot. n= represents the number of biological replicates per each organ. The green dots represent DE lncRNAs with FDR < 0.05 and log_2_ FoldChange > 2 or < -2, the blue dots represent DE lncRNAs with FDR < 0.05 and log2 FoldChange < 2 or > -2.

LncRNA expression changes in the different organs were analyzed by differential expression analysis, considering only the organs for which biological replicates were available (n ≥ 2). A series of pairwise comparisons, where the shoot organ was set as the control, were performed in both species. Overall, 1,660 and 6,396 lncRNAs exhibited significant modulation (padj < 0.05) in *T. urartu* and *Ae. tauschii*, respectively (fig. 7). In particular, the pairwise comparison root *vs.* shoot in the two wild wheats leads to a similar number of differentially expressed lncRNAs (n=1,660 in *T. urartu* and n=1,664 in *Ae. tauschii*; padj < 0.05) (supplementary file S11, S12).

To understand if the differentially expressed lncRNAs were shared among different organs, or if they were peculiar to any specific organ, a Venn diagram was created with the lncRNAs emerged as differentially expressed from each pairwise comparison (root *vs.* shoot, n=1664; seed *vs.* shoot, n=1,967; spike *vs.* shoot, n=1,729; seedling *vs.* shoot; n=1,034) in *Ae. tauschii* (supplementary fig. S4) (supplementary file S12-15). This analysis confirmed that most lncRNAs are organ-specific and, despite their low expression level, they are clearly differently modulated, especially in the most divergent organs (root, seed and spike).

## 3. Discussion

*T. urartu* and *Ae. tauschii* are the progenitors of modern wheat and the donors of the A and D genome in *T. aestivum*, respectively. The genomic tools currently available in these two diploid wild relatives represent an important opportunity to identify novel genes/alleles relevant for adaptation and consequently breeding. In fact, the diploid *T. urartu* and *Ae. tauschii* genomes are a resource of traits associated with rusticity and resistance to pathogens that have been lost during domestication. In this context the production of synthetic hexaploid wheat (*Triticum turgidum* × *Aegilops tauschii*) represents a clear example of exploitation of the genetic diversity associated with goatgrass (*Ae. tauschii*) to improve modern wheat; i.e. grain yield (Jafarzadeh et al. 2016). These advanced breeding strategies, coupled with the benefit derived from the wheat genomics era (Uauy 2017), are essential requirements for ensuring a rapid return to the applied wheat research programs, which cannot do without the deep characterization of the functional intergenic space of the genome.

In fact, although the pervasive nature of transcription of the complex genome has been proved and accepted for almost a decade (Dinger et al. 2009), the genomics studies carried out so far in plants have neglected the noncoding space, despite the fact that association studies (GWAS) have shown that the majority of GWAS hits fall outside of annotated genes and in particular within putative intergenic regulatory elements. In *T. urartu* and *Ae. tauschii* the few works have been focused on the regulatory noncoding RNAs rather than on MIR genes (Akpinar and Budak 2016; Alptekin and Budak 2016; Cagirici et al. 2017), excluding a significant portion of the genome marked as functional noncoding space (Ard et al. 2017).

Based on our extensive bioinformatics analyses, we produced the first comprehensive lncRNA repertoire of *T. urartu* and *Ae. tauschii* by leveraging public depth RNA-seq data (more than four billion Illumina reads), which enabled the detection of low expressed transcripts. This atlas of high-confidence noncoding transcripts completes the recent update of the *Aegilops tauschii* genome (Luo et al. 2017; Zhao et al. 2017), providing a detailed annotation of the functional intergenic space. This will be a fundamental integration with future efforts to improve the structure and function of the *T. urartu* genome.

More specifically, we used a genome-wide lncRNA discovery approach, applying a reference guide method coupled with an in house-developed lncRNA pipeline (De Quattro et al. 2017; De Quattro et al. 2018). This method led to the annotation of 14,131 long intergenic RNAs in *T. urartu* and 20,523 in *Ae. tauschii*, on the basis of their reference genome. Once the updated reference genome assembly (Aet v4.0) became available (Luo et al. 2017), we refined the Aet-lncRNA set by retrieving 69% of the lincRNAs previously detected in the draft genome, highlighting 312 dispersed duplicated Aet-lncRNAs. These data are in sharp contrast with the protein-coding genes in *Ae. Tauschii*, where single copy protein-coding genes account for only 13% of the total (Luo et al. 2017). This opposite trend of Aet-lncRNAs could be explained by the evidence in wheat that the rapid turnover of the intergenic space accelerates the duplication rate by 20-fold, resulting in the evolution of new functional elements (Akhunov et al. 2007). Notably, Zhao et al (Zhao et al. 2017) annotated 25,893 pseudogenes in *Ae. tauschii.* In fact, both processed and unprocessed (those that no longer have a recognizable parental gene) pseudogenes, when transcribed, can be functional through the production of lncRNAs (Milligan and Lipovich 2014).

Transposable elements have largely contributed to genome size, chromosome architecture and gene regulation of the eukaryotes genome through exaptation (Medstrand et al. 2005; Rebeiz and Tsiantis 2017). We found in both wheat progenitors that one third of lncRNA exonic sequences contains a TE domain either of *Gypsy*, *Copia* or CACTA DNA origin and less than 10% are transcribed from TE loci. In *Ae. tauschii* these three superfamilies are the most abundant, accounting for about three-quarters of the total repetitive DNA. Recently the hypothesis has been put forward that TEs may act as lncRNA functional domains and recognition sites for DNA, RNA and proteins, thus conferring them the ability to interact with and regulate other molecules (Johnson and Guigó 2014). We found a similar trend in *Brachypodium distachyon* (De Quattro et al. 2017), although in this species transposons and retrotransposons account for less than 30% of the genome size (International Brachypodium Initiative 2010). In maize the repetitive component of the genome is estimated to be 85% and it has been shown that *Copia* and *Gypsy* are the most common TE superfamilies associated with lncRNAs (Wang et al. 2017). Differently, rice lncRNAs harbor mostly miniature inverted-repeat transposable elements (MITEs) (Lu et al. 2012). This correlates with the amplification burst of LTRs and MITEs in the *Poaceae* family, which played such an important role in genome evolution. Notably, a recent study on Brassicaceae did not show an evident correlation between TE and lncRNA specificity, suggesting that this may be due to the paucity of TEs in *Brassicaceae* genomes (Nelson et al. 2016).

In our study, the lncRNAs that emerged from the association analysis with TEs appear to be highly organ-specific in both *T. Urartu* and *Ae. tauschii*, with a predominant accumulation in the young and immature spike, stamen and pistil. This suggests their involvement during the reproductive processes. In plants and animals overwhelming evidence pinpoints the expression of lncRNAs in reproductive organs, where they may exert the function of master regulators of gene expression, so coordinating the intricate molecular network activated during the reproduction phases (Golicz et al. 2018).

Despite the rapid evolution characteristic of lncRNA, a comparative genomics approach was used in this work to evaluate the degree of conservation between the wheat progenitors *T. urartu* and *Ae. tauschii* and across the temporal scale of wheat domestication and evolution by comparing the lncRNAs annotated in *T. urartu* with the subgenome A of emmer and bread wheat. Analogously, the lncRNAs from *Ae. tauschii* were compared with the subgenome D of bread wheat.

The three donors of modern wheat genomes diverged around 3 MYA, and *Ae. tauschii* is the last one that diverged, around 0.18 MYA (Middleton et al. 2014). The origin of bread wheat is dated to less than 0.4 MYA, by means of the allopolyploidization between *Triticum turgidum* (BBA^u^A^u^) and *Ae. tauschii* (DD) (Marcussen et al. 2014). This positions *Ae. tauschii* and the subgenome D of *Triticum aestivum* (bread wheat) closer within the temporal scale of wheat evolution.

Our analysis suggests a general low interspecific conservation at primary sequence level, between the two wild relatives and the species belonging to the pre-domestication and the post-domestication periods. We showed that in accordance with wheat evolution the degree of sequence conservation of the lncRNAs varies significantly between A^u^ and the subgenomes A of emmer and bread wheat compared with the *Ae. tauschii* and the subgenome D of bread wheat. This trend was also observed for the proximal *cis*-regulatory regions of lncRNAs.

Considering only the A^u^ genome, we found 557 lncRNA sequences and 1,888 regulatory regions conserved among the three species, suggesting that these lncRNAs were inherited from the wild progenitor and then stabilized in the *Triticum aestivum* after multiple rounds of hybrid speciation. Despite the close evolutionary distance that characterizes cultivated wheat and its ancestors, our results are in contrast with Clavijo et al. (2017) and show a stronger lncRNA species-specificity in wheat compared to that reported in vertebrates, i.e. 20% of inter-species conservation between human and mouse and 5% between human and fish (Ulitsky 2016), thus sustaining the hypothesis of a rapid and large sequence evolution of lncRNAs that seems to be exacerbated in wheat species. These events are likely to shape the genome organization, leading to chromosomal translocation and entire chromosome or chromosome fragment loss (Moghe and Shiu 2014). The best example of evolutionary history based on this kind of event is represented by wheat that speciated through allopolyploidy. The poor conservation that characterizes our lncRNAs, i.e. the presence in one species and the absence in other ones, could be caused precisely by the loss of specific fragments. In support of this hypothesis, a study of Feldman (Feldman et al. 1997) highlighted the elimination of specific, low-copy, noncoding sequences as a consequence of allopolyploidization. This mechanism could be one way through which the plant accentuates the difference among homoeologous chromosomes and thus ensuring an exclusive bivalent pairing of homologous chromosomes, preventing intergenomic pairing and ensuring proper chromosome segregation and, hence, full fertility (Feldman et al. 1997).

## 4. Materials and Methods

### Experimental datasets

Sixty-eight public RNA-seq libraries generated from *T. urartu* and *Ae. tauschii* were retrieved from public databases. More specifically, twelve *T. urartu* libraries produced from roots and shoots (Krasileva et al. 2013; Leach et al. 2014) were downloaded from the European Nucleotide Archive (http://www.ebi.ac.uk/ena), 15 libraries produced during the *T. urartu* genome sequencing project (Ling et al. 2013) from 7 distinct organs/conditions, i.e. leaf, root, seedling, leaf grown in control conditions, leaf grown in cold stress, shoot and spike, were downloaded from the *GigaDB* (http://gigadb.org/site/index) (supplementary Table S1). As for *Ae. tauschii*, 15 libraries from 5 organs, i.e. (shoot, root, seedling, spike and seed (Leach et al. 2014; A. Li et al. 2014) were downloaded from the European Nucleotide Archive (http://www.ebi.ac.uk/ena). From the *Ae. tauschii* genome sequencing project, 24 libraries from several organs, i.e. leaf, root, sheath, stem, spike, seed, stamen and pistil were downloaded from the *GigaDB* (http://gigadb.org/site/index) (supplementary Table S1). All the 66 libraries were sequenced as paired ends, except those from Li *et al.* (2014), which were sequenced as single end according to the Illumina technology and ranged in size between 49 and 101 bp. A total of 2,281,187,162 and 2,578,640,385 reads constituted the working sequencing dataset of *T. urartu* and *Ae. tauschii*, respectively.

The *Aegilops tauschii* genome At_v.4.0, organized in pseudomolecules together with the high-confidence gene models and TE annotation, was retrieved from http://aegilops.wheat.ucdavis.edu/ATGSP/data.php

### Bioinformatic analysis

The quality of the row reads was assessed using FastQC (Andrews et al. 2014). Then, for the libraries containing adapter contaminants, Cutadapt (v1.2.1) (Martin 2011) was applied. Quality filtered and trimmed reads were aligned to their own reference genome downloaded from Ensembl Genomes (Kersey et al. 2016): *T. urartu* version GCA_000347455.1.29, *Ae. tauschii* version GCA_000347335.1.29. Two mapping iterations were applied, with the spliced read aligner TopHat2 v2.0.9 (Trapnell et al. 2012) with an allowed number of mismatches = 0. This approach, first proposed by Cabili *et al.* (Cabili et al. 2011), maximizes the mapping efficiency by using the splice site information derived from the first mapping iteration in all samples. Successively, for each experiment we assembled the transcriptome using Cufflinks v2.0.9 (Trapnell et al. 2010) supplying the reference annotation GFF files (version Ensembl GCA_000347455.1.29 for *T. urartu* and GCA_000347335.1.29 for *Ae. tauschii*) to guide the assembly. All the transcripts assembled with Cufflinks were then merged within each species, obtaining one pool of non-redundant transcripts for *T. urartu* and one for *Ae. tauschii.* Finally, gffread, within the Cufflinks program, was used to generate a multi FASTA file, containing both novel and previously annotated transcripts.

### LncRNA identification

In order to exclude any known protein domains, we used Pfam database (Finn et al. 2014) version 27 (BlastX with E-value ≦ 0.001) to eliminate transcripts harboring any known protein domains.

The remaining transcripts were evaluated using the program Coding Potential Calculator (CPC) (Kong et al. 2007) for the quality, completeness and homology of their ORFs, and successively were compared to a database of structural RNAs derived from Rfam 12.0, representing plant species belonging to the *Poaceae* family and containing a total of 58,480 sequences (Griffiths-Jones et al. 2003). Transcripts passing all the filters were then defined as *bona fide* lncRNAs.

In order to reduce the complexity of the datasets, without losing low abundance lncRNAs, lncRNA transcripts with an overall expression of less than 10 raw read counts in total across the different organs were discarded. Read counts were calculated using the program HTSeq (Anders et al. 2014), setting no strand specificity and “intersection-noempty” so as to handle reads overlapping more than one feature. LncRNA splicing forms within each species were removed using the program CD-HIT-EST (Li and Godzik 2006), setting the sequence identity threshold at 95%. *Bona fide* lncRNAs were divided according to their genomic position in long intergenic noncoding RNAs (lincRNAs) using the Bioconductor package GenomicFeatures (Lawrence et al. 2013). Since the sequenced genomes of both *T. urartu* and *Ae. tauschii* are not assembled in chromosomes, they contain about a thousand short fragments not assembled in scaffolds, in which we identified some lncRNAs: those were not taken into account for lincRNA annotation.

### Transposable element content analysis

The annotation of TE repeats in *T. urartu* and in *Ae. tauschii* was conducted by Ling *et al.* (Ling et al. 2013) and Jia *et al.* (Jia et al. 2013) respectively. The TE information (version Ensembl GCA_000347455.1.29 for *T. urartu* and GCA_000347335.1.29 for *Ae. tauschii*) was downloaded from Ensembl DB (http://plants.ensembl.org/) and only the TEs classified according to Wicker et al (Wicker et al. 2007), for a total of 1,647,579 sequences in *T. urartu* and 1,593,537 in *Ae. Tauschii*, were considered. In addition, we also integrated this analysis with the new TE annotation provided in the new *Ae. tauschii* assembly (http://aegilops.wheat.ucdavis.edu/ATGSP/data.php).

To find the lncRNAs transcribed from an annotated TE locus, we intersected the genomic coordinates of TEs with those of lncRNAs using the Bioconductor R package *Genomic Features* (Lawrence et al. 2013), retaining only the lncRNAs fully contained within the genomic TE coordinates or vice versa. Moreover, to identify TE domains within the exonic sequences of lncRNAs, we masked the *bona*-*fide* lncRNA sequences of both species with the RepeatMasker software (Tarailo-Graovac and Chen 2009) using a custom library of TEs for every species on the basis of the above-mentioned TE assembly versions.

### EST databases to validate lncRNA transcript reconstruction

LncRNA transcript reconstruction was validated using a set of 1,327,527 *T. aestivum* ESTs downloaded from the NCBI EST database (http://www.ncbi.nlm.nih.gov/). We also took advantage of the dataset of annotated *T. urartu* noncoding transcripts from Krasileva *et al.* (2013), who reconstructed the transcriptome using a *de novo* assembly approach. Sequence similarity within our predictions and the external databases was searched using the program CD-HIT-EST-2D (Li and Godzik 2006) with a sequence identity threshold of 80%.

### Sequence conservation analysis

The level of conservation between *T. urartu* and *Ae.tauschii* lncRNAs was assessed using BLASTn (Camacho et al. 2009) with an E-value cut-off of 1e-10 and the program CD-HIT-EST-2D (Li and Godzik 2006) and a sequence identity threshold ≥ 95%. From the BLASTn output, the lncRNAs that showed a conserved multi-exon gene structure were further analyzed to assess the presence of conserved exonic splicing enhancers (ESEs) using the software SEE ESE (http://ccb.jhu.edu/software/SeeEse/index.shtml). Among the candidate ESE hexamer sequences, we retained only those that overlapped at least one of the three sequences found as very effective ESE in plants (GAAGAAGAA, CGATCAACG and TGCTGCTGG). Sequence conservation between the wild wheat relative lncRNAs and the *Triticum aestivum* and *Triticum turgidum* ssp. *dicoccoides* genomes was assessed using the BLAT algorithm with default parameters (Kent 2002). We searched each *T. urartu* lncRNA against the A genomes of *T. aestivum* and *Triticum turgidum* ssp. *dicoccoides* and each *Ae. tauschii* lncRNA against the *T. aestivum* D genome. From the hits obtained we selected only those lncRNAs with a length approximately equal to the median size length of the annotated lncRNAs in both species (465 +/- 100 bp), and among these we retained only the sequences with a maximum gap of 30 bp and that mapped just in one region of the genome. CD-HIT-EST-2D (Li and Godzik 2006) with a sequence identity threshold of 80% was used to compare the conserved lncRNAs with two *T. aestivum* EST databases, from TriFLDB (http://trifldb.psc.riken.ip/) and NCBI (http://www.ncbi.nlm.nih.gov/), and with a set of putative *T. aestivum* lncRNAs sequences deposited in the GreeNC database (Gallart et al. 2015) (http://greenc.sciencedesigners.com/).

To assess the conservation of lncRNA promoter and regulatory regions between the wild relatives, bread wheat and wild emmer wheat, we first created a .bed file with the promoter coordinates, selecting a region of 1 kbp upstream from the first nucleotide of the lncRNAs using the flank function from the Bioconductor package GenomicFeatures (Lawrence et al. 2013). With these coordinates we extracted the promoter sequences from the genome files (Ensembl *T. urartu* GCA_000347455.1.29, *Ae. tauschii* GCA_000347335.1.29 and *Triticum turgidum* ssp. *dicoccoides* 151210_zavitan_v2_pseudomolecules, *T. aestivum* IWGSC1.0+popseq.30) using BEDTools v2.17.0 (Quinlan and Hall 2010). We compared the promoter sequences from *T. urartu* with the A genomes of *T. aestivum* and *Triticum turgidum* ssp. *dicoccoides* and the promoter sequences from *Ae. tauschii* with the D genome of *T. aestivum*, using BLASTn (Camacho et al. 2009) with an E-value cut-off of 1e-10. We finally processed the BLASTn output by retaining only the sequences that matched on the reference *T. aestivum* and *Triticum turgidum* ssp. *dicoccoides* genomes with a maximum mismatch of 30 bp.

### Expression profiles and differential expression analysis

Raw counts were calculated using HTSeq (Anders et al. 2014). Expression data were normalized and expressed in Reads Per Kilobase per Million mapped reads (RPKM), using the rpkm function written in the Bioconductor package edgeR (Robinson et al. 2010). The Shannon Entropy of every transcript in the different organs was calculated using the Bioconductor package BioQC (Zhang and from Laura Badi 2015) through the function entropySpecificity (matrix, norm = T), where the matrix contained the RPKM expression values for every lncRNA in the different tissues and norm = T normalizes the scores between 0 and 1. The differential expression analysis was conducted using the Bioconductor package DESeq2 (Love et al. 2014). The following series of pairwise comparisons were taken into consideration during the analysis: root vs. shoot in *T. urartu* and root, seed, seedling and spike vs. shoot in *Ae. tauschii.* LncRNAs with adjusted p-value (FDR) ≤ 5% were considered to be differentially expressed.

## Data

Supplementary Files are available on Figshare through the link: https://figshare.com/s/73bd25fcc8326a5d38c5

## Acknowledgements

This work was supported by the International Doctoral Program in Agrobiodiversity - Scuola Superiore Sant’Anna (www.santannapisa.it).

## 6. Supplementary Material

**supplementary Table S1**

Supplemental Table S1 contains the detailed description of the public RNA-seq libraries used in this study.

**Supplementary Table S2**

supplemental table S2 contains the statistics related to raw reads, trimmed reads and alignment process.

**Supplementary Figure S1**

Schematic diagram of the bioinformatic pipeline for *T. urartu* and *Ae. tauschii* lncRNA identification.

See main text and Materials and Methods for details. In the second, dark gray box the filters for lncRNA identification are represented. The blue and violet boxes show the number of *bona*-*fide* lncRNAs annotated for *T. urartu* and *Ae. tauschii*, respectively.

**Supplementary figure S2**

Duplicated *Ae. tauschii* lncRNA

Expression level of the duplicated *Ae. tauschii* lncRNAs.

**Supplementary Figure S3**

**Shannon entropy distribution**.

Density plot showing the distribution of Shannon entropy values (X axis) for each *T. urartu* and *Ae. tauschii* lncRNA across different organs. The highest value is 1.0 and indicates specificity for one organ.

**Supplementary figure S4**

**Venn diagram of *Ae. tauschii* differentially expressed lncRNAs**.

Venn diagram representing the number of differentially expressed lncRNAs detected in every organ.

